# Long-term balancing selection in *LAD1* maintains a missense trans-species polymorphism in humans, chimpanzees and bonobos

**DOI:** 10.1101/006684

**Authors:** João C. Teixeira, Cesare de Filippo, Antje Weihmann, Juan R. Meneu, Fernando Racimo, Michael Dannemann, Birgit Nickel, Anne Fischer, Michel Halbwax, Claudine Andre, Rebeca Atencia, Matthias Meyer, Genís Parra, Svante Pääbo, Aida M. Andrés

## Abstract

Balancing selection maintains advantageous genetic and phenotypic diversity in populations. When selection acts for long evolutionary periods selected polymorphisms may survive species splits and segregate in present-day populations of different species. Here, we investigate the role of long-term balancing selection in the evolution of protein-coding sequences in the *Homo-Pan* clade. We sequenced the exome of 20 humans, 20 chimpanzees and 20 bonobos and detected eight coding trans-species polymorphisms (trSNPs) that are shared among the three species and have segregated for approximately 14 million years of independent evolution. While the majority of these trSNPs were found in three genes of the MHC cluster, we also uncovered one coding trSNP (rs12088790) in the gene *LAD1*. All these trSNPs show clustering of sequences by allele rather than by species and also exhibit other signatures of long-term balancing selection, such as segregating at intermediate frequency and lying in a locus with high genetic diversity. Here we focus on the trSNP in *LAD1*, a gene that encodes for Ladinin-1, a collagenous anchoring filament protein of basement membrane that is responsible for maintaining cohesion at the dermal-epidermal junction; the gene is also an autoantigen responsible for linear IgA disease. This trSNP results in a missense change (Leucine257Proline) and, besides altering the protein sequence, is associated with changes in gene expression of *LAD1*.

## Introduction

Balancing selection maintains advantageous polymorphisms in populations, preventing fixation of alleles by drift and increasing genetic diversity (Charlesworth 2006; Andrés 2011; Key et al. 2014). There are a variety of mechanisms through which balancing selection can act, including overdominance or heterozygote advantage (Allison 1956; Pasvol, Weatherall, Wilson 1978), frequency-dependent selection and rare-allele advantage (Wright 1939; Gigord, Macnair, Smithson 2001), temporal and spatial variation in selective pressures (Gillespie 1978; Muehlenbachs et al. 2008), or pleiotropy (Gendzekhadze et al. 2009).

When balancing selection acts on a variant long enough it creates long local genealogies, with unusually old coalescence times. Selected alleles can segregate for millions of years, with neutral diversity accumulating near the selected variant(s) due to linkage (Charlesworth, Nordborg, Charlesworth 1997; Clark 1997; Charlesworth 2006). Selection maintains alleles close to the frequency equilibrium, the frequency that maximizes fitness in the population. This results in an enrichment of variants close to the frequency equilibrium in selected and linked variation (Hudson, Kaplan 1988; Takahata, Nei 1990; Charlesworth, Nordborg, Charlesworth 1997; Charlesworth 2006). Recombination restricts these signatures to short genomic segments (Wiuf et al. 2004; Charlesworth 2006; Ségurel et al. 2012; Leffler et al. 2013). If selection is strong and constant enough, the polymorphism may survive the split of different species and persist in present-day populations of more than one species, resulting in a trans-species polymorphism (trSNP) (Muirhead, Glass, Slatkin 2002; Charlesworth 2006; Andrés 2011) (Figure 1). In species with old enough divergence time trans-species polymorphisms are rare under neutrality and are hallmarks of balancing selection (Charlesworth, Nordborg, Charlesworth 1997; Clark 1997; Wiuf et al. 2004).

**Figure 1:**
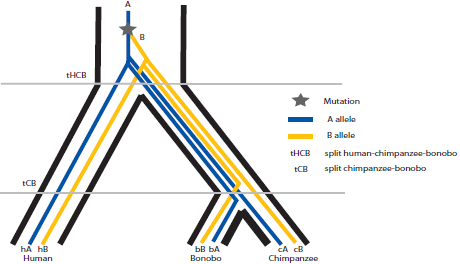
Schematic representation of a possible genealogy leading to a trans-species polymorphism (trSNP) inhuman, chimpanzee and bonobo.

The assumption that trans-species polymorphisms are very rare in humans combined with the absence of unbiased genome-wide polymorphism datasets in other great ape species resulted in few trans-species polymorphisms being described in humans: Several SNPs in the major histocompatibility locus (MHC) (Klein et al. 1993; Asthana, Schmidt, Sunyaev 2005), and a few non-MHC genes (e.g. *TRIM5* (Cagliani et al. 2010), *ZC3HAV1* (Cagliani et al. 2012), and *ABO* (Ségurel et al. 2012)). Recently, six well-defined short trans-species haplotypes containing at least two trSNPs shared in humans and chimpanzees have been identified (Leffler et al. 2013). Interestingly, none of these haplotypes contains coding SNPs, and the authors propose a role in the regulation of genes for the maintenance of these SNPs. Leffler et al. (2013) also identified a number of coding SNPs shared between humans and chimpanzees, but because filtering on allelic trees or CpG sites was not performed, it is unclear whether they represent trans-species polymorphisms or recurrent mutations (an important question in the identification of trSNPs, see below).

Here we analyze the exomes of 20 humans, 20 chimpanzees and 20 bonobos to identify trans-species polymorphisms present since the *Homo-Pan* common ancestor until the present-day population of each of the three species. By including the three species we focus only on strong balancing selection that has been maintained in the three lineages. Besides identifying coding trSNPs in several MHC genes, we also identify a novel trans-species polymorphism (rs12088790) maintained by long-term balancing selection in the gene *LAD1 (ladinin-1)*.

## Results

### A model for neutral trSNPs in humans, chimpanzees and bonobos

As mentioned above, the presence of neutral trSNPs is unlikely when species diverged long ago. To estimate how probable a shared SNP would be in a sample of SNPs from one of the three species, we developed a model based on coalescent theory (Supplementary Information I), assuming the ancestral and the species-specific population sizes estimated in Prado-Martinez et al. (2013). Under this model, given that the lineages of bonobos and chimpanzees diverged only about 2 million years ago (Prüfer et al. 2012) and their present-day populations share polymorphisms, we expect, under neutrality, 0.85% of the SNPs in bonobos to be segregating in chimpanzees, and 4.6% of chimpanzee SNPs to also be segregating in bonobos (see Supplementary Information I). Conversely, a neutral trans-species polymorphism between *Homo* and any of the two *Pan* species is unlikely to occur by genetic drift alone: We estimate that a SNP found in a sample of humans has a probability P_HC_ = 1.6×10^−8^ of being polymorphic in chimpanzees too (see also Supplementary Information I). The model also allows us to calculate the probability of observing a SNP shared by all three species (bonobo, chimpanzee and human) in a sample of human SNPs. This probability (called P_FINAL_) is, under neutrality, approximately equal to 4.0×10^−10^. This is roughly 39 times lower than the probability that a SNP in humans is also polymorphic in chimpanzees (P_HC_), illustrating the advantage of including bonobos in the comparison. Given that we observe 121,904 human SNPs, we expect about 5.0×10^−5^ neutral trSNPs in the three species. We note that these are actually overestimates, since coding variation is subject to purifying and background selection that produce shallower coalescent trees than neutrally evolving loci. Therefore, any trSNP that we find is highly unlikely to have occurred under neutrality. An exploration of the behavior of the model under a range of parameters for the split times and population sizes is detailed in Supplementary Information I. We note that the parameters needed to explain the presence of neutral trSNPs in the three species are unrealistic, given our knowledge of human and great ape demographic history.

### Idenfication of trSNPs

We sequenced the exomes of 20 Yoruba humans, 20 central chimpanzees *(Pan troglodytes troglodytes)* and 20 bonobos *(Pan paniscus)* to an average coverage of ~18X in each individual (data is very homogeneous across species in coverage and quality, see Materials and Methods). We uncovered a total of 121,904 high-quality SNPs in human, 262,960 in chimpanzee and 99,142 in bonobo. This represents a novel SNP discovery rate of ~33.54% in bonobo, ~49.29% in chimpanzee and ~2.8% in human (compared with Prado-Martinez et al. (2013) and dbSNP build 138). We focused on the 202 coding SNPs with the same two segregating alleles in the three species, the *shared SNPs* (shSNPs).

Two important confounding factors in the identification of trSNPs are genotype errors and recurrent mutations. To limit the influence of genotype errors in the form of mapping and sequencing artifacts, we conservatively removed SNPs that: 1) are in the upper 5% tail of the empirical distribution of coverage in at least one species; and 2) do not lie in regions of unique mappability of 24mer. We further removed SNPs that are not in Hardy-Weinberg equilibrium (HWE) with p-value (*p*) < 0.05 in at least one species (see Supplementary Information II). Regarding recurrent mutations, they are particularly likely in hypermutable sites where the probability of a parallel mutation in two lineages is high. Examples of this are CpG dinucleotides (where a methylated cytosine can deaminate to a thymine and result in a C–>T transition (Bird 1980; Hodgkinson, Eyre-Walker 2011), but additional, cryptic heterogeneity in mutation rate exists (Hodgkinson, Ladoukakis, Eyre-Walker 2009; Hodgkinson, Eyre-Walker 2010; Johnson, Hellmann 2011).

Removing CpGs could reduce the number of recurrent mutations, but SNPs associated with CpGs represent a large fraction of SNPs in the genome (about 25% of human SNPs) and recurrent mutations can also occur at non-CpG sites (Hodgkinson, Eyre-Walker 2011). We therefore mark but consider CpG SNPs, and use additional lines of information to tell apart trSNPs from recurrent mutations. Specifically, SNPs that result from recurrent mutations are expected to fall in genomic regions that follow the species tree (Figure 2)because the most recent common ancestor of the genomic segment containing a human SNP falls in the human branch, predating (backwards in time) the coalescence of lineages from the different species (see previous section). On the contrary, trans-species polymorphisms create local genealogies that cluster by allele (Figure 2) because the most recent common ancestor of the genomic segment containing the trSNP predates the split of the three species (Schierup, Mikkelsen, Hein 2001; Wiuf et al. 2004). Therefore, a SNP’s surrounding region allows us to distinguish trSNPs from recurrent mutations.

**Figure 2:**
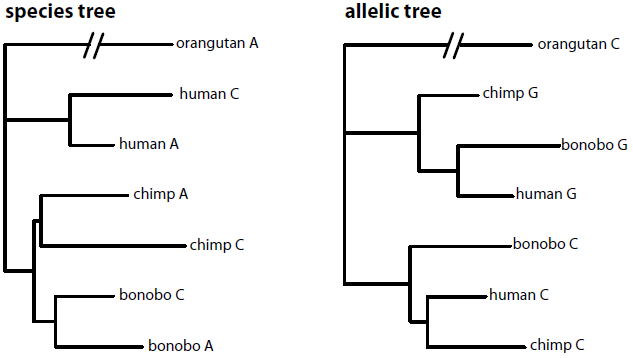
Examples of a species tree and an allelic tree using sixhaplotypes, one per species and allele. Each Neighbor-joining tree is computed on a 500 bp region around a shSNP in our dataset for the genes *TXNDC2* (species trees) and *HLA-DQA1* (allelic tree).

For each shSNP we inferred the phylogeny of its genomic region (Materials and Methods) and considered further only shSNPs that fall in genomic regions that exhibit trees that cluster by allelic type. Of the 202 original shSNPs, and after additional filtering (coverage, mappability and HWE), only 20 have a probability of an allelic tree (P_allelic_) > 0.90 (Table S4); these shSNPs, all of which are present in dbSNP build 138, were considered ‘candidate trSNPs’. They lie in 15 different genes, including three HLA genes. Figure 3 shows the neighbor-joining tree of one of such trSNPs, the one present in gene *LAD1,* with sequences clustering by allelic type. Only two ‘candidate trSNPs’ (both in *HLA-DQA1)* are not associated with CpG sites (Table S4). We also note that other HLA genes that have been described before as being targets of balancing selection in humans have shSNPs that were excluded from our analysis due to the stringent filtering criteria implemented, although no specific filters were applied in the MHC region.

**Figure 3:**
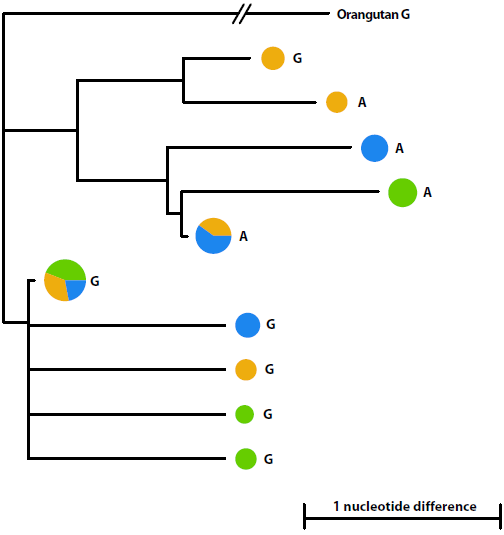
Neighbor-joining tree of *LAD1*. The tree was constructed using a 350 bp region as described in Methods. The size of the pie charts is proportional to the number of haplotypes (n=120), with colors representing the species. The alleles of the trSNP are shown next to the pie charts. The orangutan sequence (PonAbe2) was used as outgroup. Three chimpanzee haplotypes carrying the G allele cluster with haplotypes carrying the Aallele, likely due to a recombination event (more likely to occur in chimpanzee, the species with the largest effective population size).

Because trSNPs have been previously described in HLA genes (Lawlor et al. 1988; Mayer et al. 1988; Fan et al. 1989; Klein et al. 1993; Asthana, Schmidt, Sunyaev 2005; Leffler et al. 2013) we focus on the remaining genes (13 ‘candidate trSNPs’). Our filtering criteria exclude the majority of systematic sequencing errors, so we next investigated the possibility of mapping errors due to collapsed paralogs (when paralogs are very similar in sequence, mapping errors can result in erroneous SNP calls). We BLAT (Kent 2002) the 25 bp region surrounding the 13 non-HLA candidate trSNPs to the reference genome sequences of human (hg19) and chimpanzee (PanTro4). Only four ‘candidate trSNPs’ (in genes *LY9, LAD1, SLCO1A2* and *OAS1)* map uniquely to all genomes, with the remaining nine candidate trSNPs mapping to regions that have a close paralog in at least one species (see Supplementary Information II and Table S4). Although this does not discard these positions as SNPs in the other species (or in the species with non-unique BLAT hits) we conservatively removed them from further analyses. We therefore focus on these four SNPs, to investigate additional signatures of long-term balancing selection.

### The probability of an allelic tree

As the allelic tree provides very strong evidence for a SNP to be a trSNP, we next aim to determine how likely an allelic tree is, for each ‘candidate trSNP’, under recurrent mutation. To answer this question we ask how often we observe an allelic tree of the same length and minimum number of informative sites as those of the ‘candidate trSNPs’. We estimated the false discovery rate (FDR, the chance of obtaining an allelic tree under recurrent mutation) for each allelic tree length by analyzing random SNPs in the genome of the three species. In short, we pair random SNPs in the human genome with a close-by SNP in chimpanzees and bonobos; these nearby, independent mutations act as pseudo-recurrent mutations where to investigate the neutral probability of an allelic tree (see Materials and Methods, SuppIementary Information III and Table S1 for details). We found that, as expected, the FDR is inversely proportional to the length of an allelic tree (Supplementary Information III); that is, the longer the genomic region, the lower the FDR of an allelic tree because the number of phylogenetically informative positions grows and so does the chance for recombination. If we condition on observing additional informative sites (besides the ‘candidate trSNP’) the FDR drops substantially and becomes more uniform across the different lengths (Table S1).

For our set of candidate trSNPs, and after considering the exact number of informative sites uncovered in the length of each allelic tree, only *LY9*’s allelic tree shows high FDR (36.8% for 100bp and 4 informative sites). All other candidate trSNPs fall in allelic trees that given the number of informative sites uncovered in each tree have low FDR (Table 1).

**Table 1:**
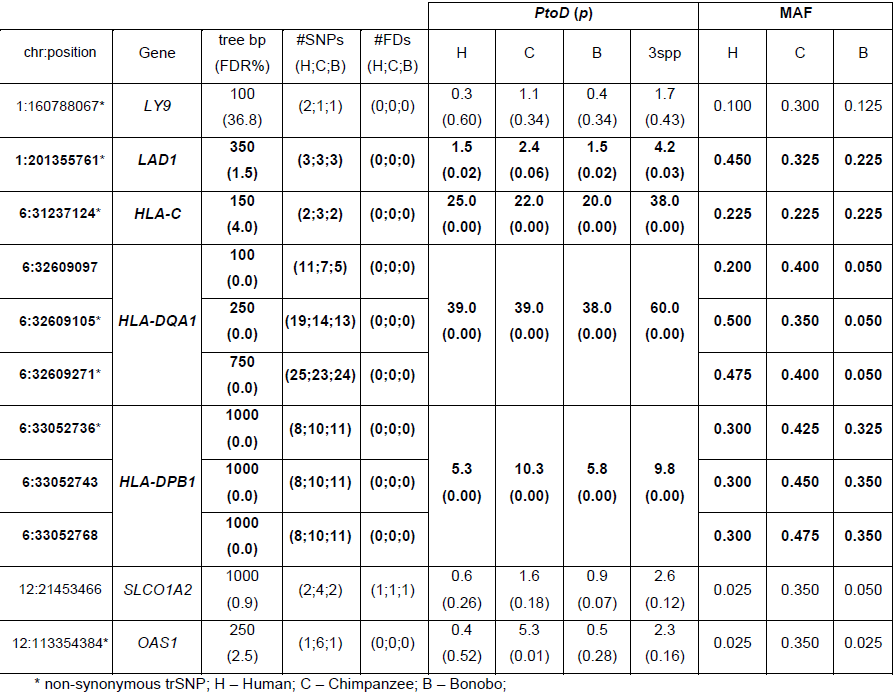
Comparison of different signatures in candidate trSNPs and genes. The estimated length of the allelic trees (and respective FDR), the number of polymorphisms (#SNPs) and fixed differences (#FDs) in the allelic tree, the polymorphism-to-divergence *(PtoD)* ratios for the whole gene and minor allele frequencies (MAF) of trSNPs are shown. For ‘Human’, we present the *PtoD* ratio obtained in the human-bonobo comparison, which is very similar to the human-chimpanzee comparison. The genes with trSNPs and consistent signatures of long-term balancing selection are shown in bold.

### Excess of polymorphism linked to the trSNPs

We further investigate whether, as expected under long-term balancing selection, the ‘candidate trSNPs’ fall in regions that exhibit an excess of genetic diversity after taking heterogeneity in mutation rate into account. We calculated the ratio of polymorphism to divergence *(PtoD = p/(d+1)*, where *p* is the number of polymorphisms identified in a species and *d* the number of fixed differences identified between species – see Supplementary Information IV) in the genes containing our four non-HLA ‘candidate trSNPs’ *(LY9, LAD1, SLCO1A2, OAS1);* we also analyze the seven HLA trSNPs *(HLA-C, HLA-DQA1* and *HLA-DPB1)*. For each gene we investigate different genomic regions, in each species: a) ‘ALL’ – the entire genic region; b) ‘coding’ – only their coding exonic sequence; c) ‘500bp’ – the 500 bp surrounding the trSNP; and d) the ‘length of allelic tree’ (Table S4). First, if we focus on individual genes, *HLA* genes are in the very far tail of the empirical genomic distribution of *PtoD*, with a significant excess of polymorphism in all the comparisons performed (Table S4). For non-HLA genes, only *LAD1* shows a consistent excess of diversity in the three species, with most comparisons being significant in human and bonobo, and marginally non-significant in chimpanzee (see Tables 1, 2 and S4, and Supplementary Information IV). The weaker signal in chimpanzee is likely due to this species’ larger effective population size (Prado-Martinez et al. 2013) that translates in higher genomic diversity and lower power to detect the localized increased diversity in *LAD1*. The signal is weaker for the other three genes. No excess of polymorphism is observed in *SLCO1A2,* and in *LY9* high *PtoD* values are observed only for the ‘length of tree’, due to the presence of a single additional SNP in humans (in such a small region). *OAS1* shows significant excess of polymorphism only in chimpanzee.

**Table 2:** *PtoD* ratios calculated in the gene *LAD1* (with the corresponding percentile in the empirical distribution in parenthesis). For ‘Human’, we present the *PtoD* ratio obtained in the human-bonobo comparison, which is very similar to the human-chimpanzee comparison.

|  |  | Human | Chimpanzee | Bonobo |
| --- | --- | --- | --- | --- |
| *PtoD* | ALL | 1.50 (0.023) | 2.40 (0.059) | 1.50 (0.019) |
|  | Coding | 2.00 (0.018) | 1.25 (0.332) | 1.00 (0.068) |
|  | 500bp | 2.00 (0.024) | 1.50 (0.317) | 1.50 (0.069) |
|  | Length allelic tree | 3.00 (0.028) | 3.00 (0.074) | 3.00 (0.024) |
|  | 3spp | 4.20 (0.028) |  |  |

We also calculate a three-species *PtoD* (‘3spp’) for the entire genic region by jointly considering (the union of) all polymorphisms and divergent sites across the three species. The ‘3spp’ *PtoD* is unusually high in all HLA genes (p ≤ 0.002) and in *LAD1* (p = 0.028), but not in the other three genes (Table 1, 2 and S4). In fact, only 0.005% of genes in the genome have, in each of the three species, a p-value equal or lower than that of *LAD1.* This shows that the combined excess of diversity of *LAD1* in all three species is highly unusual. In addition, we note that *LAD1’s* signature is due to the strong enrichment in polymorphism in the region surrounding the trSNP (rs12088790): All SNPs we identified in *LAD1* are within 182bp of rs12088790.

Taken together, these results indicate that apart from the three *HLA* genes, only *LAD1* has a signature of long-term balancing selection in the three species. *OAS1* shows signatures of balancing selection in central chimpanzees, which have been previously reported (Ferguson et al. 2012), but the gene shows rather unremarkable signatures in bonobo and human (Table 1). We cannot discard the possibility that *SLCO1A2, LY9* or *OAS1* have been under balancing selection, but conservatively we focus on *LAD1, HLA-C, HLA-DQA1* and *HLA-DPB1* as our final set of trSNPs.

The set of these four genes is, in all species, significantly more polymorphic than the empirical distribution of all genes with at least one variable site (polymorphism or substitution) in our dataset (Tables S2 and S3, and Figures S5 and S6). *LAD1* is the least polymorphic of the four genes, which is not surprising as the remaining trSNPs fall in *HLA* genes.

### Intermediate-allele frequency of the trSNPs and linked variants

The allele frequency distribution of sites linked to a balanced polymorphism is expected to exhibit an excess of alleles at frequencies close to the frequency equilibrium. If the frequency equilibrium is high enough (e.g. 0.5) the local site frequency spectrum (SFS) will show an observable departure from the genome-wide empirical distribution. We note that the frequency equilibrium can be at any allele frequency, so while an excess of intermediate-frequency alleles is indicative of balancing selection, this is not a necessary signature.

The SFS of the four genes together shows a significant shift towards intermediate-frequency alleles, in all species (Mann-Whitney U test *p* < 4×10^−10^; Figure 4 and (Table 3). When we consider the genes individually, almost all exhibit a significant excess of intermediate-frequency alleles in all species except for *LAD1* in bonobo and human (marginally non-significant), and for *HLA-C* in bonobo (Table 3). When we combine all SNPs in each gene (the union of SNPs in all three species) and compare the resulting SFS with the combined empirical SFS (the union of all SNPs from all three species), all genes show a significant shift towards intermediate frequencies (Mann-Whitney U test *p* ≤ 0.046, (Table 3), including *LAD1*.

**Figure 4:**
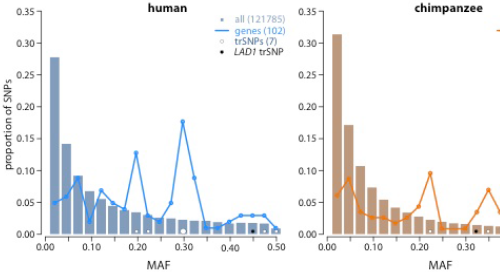
Folded site frequency spectra (SFS) of trSNPs and other SNPs in the genes. The x-axis represents the minor allele frequency (MAF) and the y-axis the proportion of sites in that frequency bin. The histograms show the spectrum of the entire exome (‘all’) for each species, excluding the four genes containing a trSNP; the lines show the combined SFS of all SNPs in the four genes containing a trSNP. The number of SNPs in each category is annotated in the legend. The trSNPs are shown as empty circles, with size proportional to the number. A black circle represents the trSNP in LAD1.

**Table 3:** P-values (Mann-Whitney U test) for excess of intermediate-frequency alleles comparing the SFS of the genes to the genome-wide SFS.

| GENE | Human | Chimpanzee | Bonobo | 3spp |
| --- | --- | --- | --- | --- |
| *LAD1* | $5.7 \times 10^{-2}$ | $4.3 \times 10^{-2}$ | $7.4 \times 10^{-1}$ | $4.6 \times 10^{-2}$ |
| *HLA-C* | $2.5 \times 10^{-2}$ | $1.8 \times 10^{-5}$ | $5.4 \times 10^{-2}$ | $3.1 \times 10^{-7}$ |
| *HLA-DQA1* | $1.4 \times 10^{-6}$ | $1.9 \times 10^{-12}$ | $4.5 \times 10^{-3}$ | $3.4 \times 10^{-19}$ |
| *HLA-DPB1* | $4.4 \times 10^{-9}$ | $5.1 \times 10^{-17}$ | $1.2 \times 10^{-11}$ | $1.7 \times 10^{-36}$ |
| all four genes | $3.9 \times 10^{-14}$ | $2.0 \times 10^{-30}$ | $3.7 \times 10^{-10}$ | $1.8 \times 10^{-54}$ |

The trSNP in *LAD1,* which is a missense polymorphism, is at intermediate frequency in all three species (Table 1) and Figure 4): MAF=0.450 in human, 0.325 in chimpanzee, and 0.225 in bonobos. These frequencies are all in the upper quartile of the empirical allele frequency distributions of non-synonymous variants: In the upper 1.9% quantile for human, in the 8.6% for chimpanzee, and in the 23.8% in bonobo.

When we investigate the 1000 Genomes dataset (Abecasis et al. 2012), which contains both coding and non-coding data for *LAD1*, we observe a significant excess of intermediate-frequency alleles in all African populations, although the signature varies across human groups with some non-African populations showing an excess of low-frequency alleles instead (Table S6). The trSNP is itself present in all these populations throughout the world: At intermediate frequency in all African populations (31% < MAF < 48%) and at lower frequency (MAF < 8%) values for *LAD1’s* trSNP between the African Yoruba and two non-African populations (Toscani and Han Chinese) we observe high allele frequency differences (F_ST_ = 0.238 and 0.293, respectively), which are in the top 6.5% tail of the empirical F_ST_ distribution. The polymorphism is thus shared across human populations, but its frequency shows high levels of population differentiation among human groups.

### Balancing Selection in *LAD1*

*LAD1 (ladinin 1)* spans 18,704 bp and is composed of 10 exons. We obtained a total of 1,213bp of the gene by sequencing the complete exons 4, 7 and 9, as well as parts of exons 2, 3 and 5. The trSNP found in *LAD1* (chr1: 201355761, rs12088790) encompasses an A/G polymorphism (reverse strand), which we validated with Sanger sequencing, and is a missense mutation located in exon 3 that results in a Leucine to Proline change. This change has a moderately conservative Grantham score (amino acid replacement score based on chemical similarity - Leucine –> Proline = 98) (Grantham 1974).

Besides altering the sequence of the protein, the trSNP is associated with expression changes in present-day humans. Specifically, when we analyzed expression data in lymphoblast cell lines from a subset of the 1000 Genomes project individuals (Lappalainen et al. 2013), we observed significantly lower expression of *LAD1* in carriers of at least one ancestral *G* allele (GG and GA genotypes) than in AA homozygotes (*p* = 0.02). Comparing carriers of at least one *A* allele with GG homozygotes did not show a significant difference in expression levels (*p* = 0.21). This shows that the derived *A* allele is associated with increased expression of *LAD1* in an at least partially recessive manner. Mapping biases are not responsible for this result as the total number of SNPs uncovered in the closest region (one additional SNP in the 150 bp region that affects read mapping) is only moderate.

## Discussion

By comparing the exomes of humans, chimpanzees and bonobos, we identify polymorphisms maintained by long-term balancing selection in the *Homo-Pan* clade. Undoubtedly, other cases of long-term balancing selection exist, including species-specific balancing selection (Pasvol, Weatherall, Wilson 1978; Bamshad et al. 2002; Wooding et al. 2004; Wooding et al. 2005; Muehlenbachs et al. 2008; Andrés et al. 2009; Andrés et al. 2010), but here we focus on selection that is old, strong, constant and shared across lineages, and that results in trans-species polymorphisms. Even among trSNPs, we focus only on coding variants shared among the three species, and likely underestimate the number of human trSNPs. First, by focusing on coding variation we are blind to balancing selection that maintains variants outside genes, which may not be rare (Leffler et al. 2013). Second, by restricting on a SNP being present in the three species we discard cases where the variant was lost in one of the lineages, which may again not be rare. Even one of the best-established cases of trSNPs, the one present in the *ABO* gene from humans to old word monkeys, is not shared among the three species because it was lost in chimpanzees (Ségurel et al. 2012). This is not unexpected as it is likely that one of the species has undergone demographic or selective changes that weakened or changed selection on an old balanced polymorphism. Conversely, considering three species *(e.g.* adding bonobo) reduces the probability of trSNPs under neutrality; in fact, after considering the number of SNPs discovered in humans (121,904), we expect to observe no neutral trSNP (specifically, we expect 5.0×10^−5^ neutral trSNPs). Consistent with this, the majority of coding shSNPs we identified are likely the result of recurrent mutations, as they fall in genomic regions whose phylogenies agree with the expected species tree.

We identify seven trSNPs that pass our filtering criteria and that cluster by allelic tree, with an extremely low probability under recurrent mutation. The loci containing these seven SNPs show, in addition, the excess of polymorphism expected under longterm balancing selection. Six trSNPs are located in *HLA* genes *(HLA-DQA1, HLA-C* and *HLA-DPB1)* and one is a non-synonymous SNP in exon 3 of the gene *LAD1* (rs12088790). This variant, which has segregated for millions of years in these lineages, represents to our knowledge the only trans-species polymorphism known to segregate in present-day populations of these three species outside of the MHC. As for the remaining candidate trSNPs, the combined results of our analyses are not strong and consistent enough to provide unequivocal evidence that these are targets of long-term balancing selection (although they can be). We thus focus on *LAD1*, where the evidence is clear.

Besides containing a trSNP whose genomic region clusters by allelic type, *LAD1* exhibits high levels of genetic diversity (particularly in bonobos and humans) and it shows excess of intermediate-frequency alleles (significant in chimpanzee and marginally non-significant in humans, although highly significant in the 1000 Genomes’ Africans). *LAD1* is thus an unusual gene in its consistent signatures of long-term balancing selection.

The trSNP, rs12088790, segregates at intermediate frequency in Yoruba, bonobos and chimpanzees. It is present in all 1000 Genomes human populations (Abecasis et al. 2012), although at intermediate frequency in African populations and at low frequency in non-African populations. It is not uncommon for targets of long-term balancing selection to show population differences in the allele frequency distribution (Andrés et al. 2009), sometimes due to changes in selective pressure across human groups (de Filippo et al., in preparation). The fact that only African populations show a significant excess of intermediate frequency alleles in *LAD1* (Table S6) and that F_ST_ for rs12088790 is high between African and non-African populations, suggest that this might be the case for *LAD1.* Although speculative, we note that it is possible that some environmental pressures long shared by humans, chimpanzees and bonobos, and that still affect certain African populations, have changed in human populations living in different environments outside of Africa.

Although rs12088790 in *LAD1* is a good candidate to have been the target of selection (being non-synonymous and present in the three species), it is possible that it is instead maintained by linkage to an undiscovered selected trSNP, as the maintenance of several linked trSNPs is possible under long-term balancing selection (Ségurel et al. 2012). Although more detailed genomic and functional analysis on *LAD1* are needed to completely clarify this question, we explored a recently published catalog of great ape genetic polymorphism in search for additional human-chimpanzee-bonobo shSNPs (Prado-Martinez et al. 2013). Besides rs12088790 (which in that dataset also segregates in all three species), we identified one additional shSNP in the three species. This SNP (rs12035254, chr1:201349024) is intronic and downstream of exon 10, and is located about 6 kbp downstream rs12088790 (see Supplementary Information VI). The distance between the two SNPs makes it unlikely that rs12035254 is responsible for the very localized signatures in rs12088790’s genomic region. We further compared the trSNPs found in this study with a list of shSNPs between human and western chimpanzee provided by Leffler et al. (2013) but were unable to retrieve them. This is likely due to different sampling and sequencing strategies adopted in the two studies (see Supplementary Information VI). Nonetheless, Leffler et al. (2013) also reported several human-chimpanzee shSNPs in the genes *HLA-DQA1* and *HLA-DPB1*, although the specific variants are different from the ones uncovered here.

Interestingly, the two alleles of rs12088790 are associated with differences in expression levels of *LAD1*, with higher expression associated with the ancestral G allele in lymphoblastoid cell lines. This highlights the possibility that, in addition to causing an amino acid replacement, the trSNP might also have regulatory effects (although we cannot discard the possibility that another, nearby variant, is responsible for the observed differences in expression).

The precise biological mechanisms leading to long-term balancing selection on *LAD1* are not known. The gene encodes a collagenous anchoring filament protein of basement membrane at the dermal-epidermal junction. The mRNA and the protein are observed in a number of tissues including the gastrointestinal system (and its accessory organs), the kidney, prostate, placenta, and one type of hematopoietic cells (Kim et al. 2014). Genes involved in cell adhesion and extracellular matrix components are enriched among candidate targets of balancing selection and among genes with intermediate-frequency alleles in pathogen-rich environments (Andrés et al. 2009; Fumagalli et al. 2009; Fumagalli et al. 2011; Key et al. 2014). This suggests that certain components of the cellular junction may benefit from the presence of functional polymorphism, perhaps as a defense against pathogens. In this context, *LAD1* may represent one of such examples.

Interestingly, genetic variation in *LAD1* is associated with linear IgA disease, an autoimmune blistering disease. The disease, which affects mostly children and elderly adults (McKee, Calonje, Granter 2005), is caused by the presence of circulating IgA autoantibodies that target peptides in the Ladinin-1 protein, causing an immunological reaction. This results in the disruption of the dermal-epidermal cohesion, leading to skin blistering that predominantly affects the genitalia but also the face, trunk and limbs (Ishiko et al. 1996; Marinkovich et al. 1996; Motoki et al. 1997; McKee, Calonje, Granter 2005). Although our understanding of the effect of the disease in different populations is biased by the fact that the disease (which is rare) has mostly been studied in Western countries, some evidence suggests that it is more common in Africa (Aboobaker et al. 1991; Denguezli et al. 1994; Monia et al. 2011). Balancing selection has been proposed to play a role in the evolution of autoimmune genes, because the inflammatory response must be precisely balanced to be effective yet moderate (Ferrer-Admetlla et al. 2008). Whether balancing selection in *LAD1* is responsible for its role in auto-immunity remains though unclear. It is possible, and perhaps more likely, that autoimmune diseases appear as consequences of diversity in proteins that is maintained by balancing selection and happen to be able to initiate pathogenic immunological reactions. Further work is necessary to discern the functional consequences and advantageous role of its balanced polymorphisms in humans and other primates.

## Materials and Methods

### DNA samples and sequencing

We performed whole-exome capture and high-coverage sequencing of 20 humans, 20 central chimpanzees *(Pan troglodytes troglodytes)* and 20 bonobos *(Pan paniscus)*. Human samples belong to the well-studied Yoruba population from HapMap; bonobo and chimpanzee blood samples were collected in African sanctuaries (Lola ya bonobo sanctuary in Kinshasa, Democratic Republic Congo; and Tchimpounga sanctuary, Jane Goodall Institute, Republic of Congo, respectively) and immortalized as cell culture (Fischer et al. 2011). DNA was extracted using the Gentra Purgene Tissue Kit (Qiagen), sheared to a size range of 200 to 300 bp using the Bioruptor (Diagenode) and converted into DNA libraries for capture and sequencing (Meyer, Kircher 2010). All samples were double-indexed to prevent cross-sample contamination during the processing and sequencing of the samples (Kircher, Sawyer, Meyer 2012). Exome capture was performed using the SureSelect Human All Exon 50Mb Kit (Agilent Technologies). The kit design is based on the complete annotation of coding regions from the GENCODE project with a capture size of approximately 50 Mb. We selected all Ensembl genes (mapping uniquely to hg19) that are RefSeq genes (with good functional support) and targeted by our capture design, and selected their longest RefSeq transcript. Samples were then pooled by species and sequencing was performed on Illumina’s GAIIx platform, with paired-end reads of 76bp.

### Base calling and read mapping

Base calling was performed with Ibis (Kircher, Stenzel, Kelso 2009), and reads with more than 5 bases with a base quality score lower than 15 were discarded. Reads were aligned to the human reference genome hg19 using BWA with default parameters. Mapping all individuals to the same reference genome prevented complications from mapping to genomes of different quality. Only reads with a mapping quality (MQ) ≥ 25 and mapping outside of known segmental duplications in the three species were considered for further analysis. Specifically, the average coverage for each individual is 18.9X in human, 17.9X in chimp and 17.9X in bonobo.

### Genotype calling and filtering

Genotype calls were performed in the autosomes using the Genome Analysis Toolkit (GATK) *UnifiedGenotyper* (version 1.3-14) (McKenna et al. 2010). Aside from true variation, these preliminary SNP calls likely include false positives due to the presence of mismapped reads, misaligned indels and systematic errors. We used a combination of strict filters to remove such errors. SNPs were removed using the following criteria (acronyms correspond to the GATK package or fields in the VCF files):

- The depth of coverage (DP) was <8 or >100 in at least 50% of the individuals of each species. This allowed us not only to exclude positions for which the coverage depth was low, but also positions that might fall in segmental duplications not annotated in the datasets above [28-30];
- The quality score (QUAL) of the call was <50;
- There was evidence of strand bias (SB>0);
- The genotype quality (GQ) was <10 in all individuals carrying the alternative allele;
- The SNP was located within 3bp of a homopolymer with a minimum length of 5bp;
- The SNP was located within 5bp up- and down-stream of an insertion or deletion (indel) polymorphism or substitution with the human reference genome.

### Shared SNPs as trans-species polymorphisms

Wrongly mapped reads are difficult to account for and can result in an increased false discovery of shSNPs. In order to remove undetected duplications, we further filtered shSNPs to remove sites with unusually high coverage, that are in Hardy-Weinberg disequilibrium, and that do not lie in unique regions of the genome (see Results).

### Haplotype inference and allelic trees

We use the fastPHASE 1.4.0 software (Scheet, Stephens 2006) to infer the chromosomal phase for the alleles of each of the genes containing at least one shSNP. The inferences were performed separately for each species and for each chromosome using the default parameters of fastPHASE.

The region surrounding a trans-species polymorphism is expected to follow unusual genealogies where haplotypes cluster by allelic type rather than by species. This occurs because the age of the balanced polymorphism predates the speciation time and, unless recombination happens, there will be no fixation of new mutations. We call these two types of phylogenies “allelic tree” and “species tree” (Figure 2). The trees were inferred in windows of different lengths (from 100 bp to 2,000 bp) centered on the shared polymorphism, as the region expected to follow the allelic tree is very short due to the long-term effects of recombination. We considered as candidate trans-species polymorphisms only shSNPs that show an allelic tree with probability (P_allelic_) > 0.9 in a window of at least 100 nucleotides.

We adopted a simple resampling approach to calculate P_allelic_ in the region surrounding a shSNP. We randomly created 1,000 samples of six haplotypes (one haplotype per allele and per species). For each of the 1,000 resamples we built a neighbor-joining tree using as distance matrix the number of nucleotide differences among the six haplotypes. If the three closest tips were haplotypes from the three species containing the same allele of the shSNP, it was considered an allelic tree. If the two different human haplotypes are closer to each other than to any other haplotypes, the tree was considered a species tree (the relationship between chimpanzees and bonobos was not considered because shared polymorphism can occur given their short divergence time). P_allelic_ was estimated as the proportion of resampled trees that were allelic trees. Figure 2 shows an example of allelic and species trees built from six haplotypes.

We also estimated the probability to observe an allelic tree of a given length (the false discovery rates, FDRs) under recurrent mutation and based on our empirical dataset. For each observed allelic tree lengths (Table S1), we randomly chose 1,000 human SNPs and the closest SNP in chimpanzee and bonobo. We then ‘paired’ these SNPs (i.e. use the allelic information of each SNP) as if they occurred in the same genomic position rather than in different positions, and calculated Pallelic for these haplotypes (based on the alleles found in each species’ SNP). Because these SNPs arose from independent mutations in each lineage, they perfectly mimic a recurrent mutation (falling at the same site) in the three species. The proportion of random samplings with P_allelic_ > 0.9 (i.e. the criterion used to consider the trSNP) reflects the FDR for that given length.

### Polymorphism-to-divergence ratios (PtoD)

We defined the ratio of polymorphism to divergence *PtoD = p/(d+1)*, where *p* is the number of polymorphisms observed in a species and *d* the number of fixed differences between species. For each candidate gene, we estimated significance based on the percentile of each candidate in the empirical genomic distribution of all genes.

In order to ascertain significance when comparing the set of candidate loci to the set of control loci (empirical distribution), we performed 2-tail Mann-Whitney U (MW-U) tests and used a critical value of 5%. After comparing the *PtoD* values in the two groups, we sequentially removed the top candidate gene (i.e. one gene each time) from the candidate’s group and recalculated MW-U p-values maintaining the control group unaltered (see Supplementary Information IV for details).

### Measuring expression levels in *LAD1* alleles

We analyzed lymphoblastoid cell line expression data obtained from a subset of 462 of the 1000 genomes project individuals provided by Lappalainen et al. (2013). To compute gene expression we used the aligned reads provided by Lappalainen et al. (2013) and assigned reads with a mapping quality (MQ) ≥ 30 to protein coding genes by overlapping the read coordinates with gene coordinates (ENSEMBL version 69). Reads overlapping a gene are summed up and used as the estimate for gene expression.

We grouped the individuals by their genotype at position chr1:201355761 (rs12088790, the non-synonymous trSNP in *LAD1)*. We sought to test for allele-specific expression for *LAD1* between individuals carrying the two different trSNP alleles by testing for differential expression between (i) the groups of individuals with genotype AA vs. GG/GA and (ii) the groups of individuals with genotype GG vs. GA/AA. We computed differential expression for *LAD1* for (i) and (ii) using the DESeq package (Anders, Huber 2010). Expression values in both groups are modeled by a fit of a negative binomial distribution. DESeq tests then for differences between the distributions of the two groups.

## Acknowledgments

We thank the sequencing group at the Max Planck Institute for Evolutionary Anthropology and Martin Kircher for data production. We also thank Felix M. Key, Gabriel Renaud, Jing Li, Kay Prüfer and Joshua Schraiber for helpful discussions and suggestions, as well as four anonymous reviewers, whose comments improved substantially the manuscript. We are grateful to the Lola Ya Bonobo sanctuary in Kinshasa, Democratic Republic Congo, and the Tchimpounga sanctuary, Jane Goodall Institute, Republic of Congo, for allowing access to the primate samples. This work was supported by the Max Planck Society. JCT is supported by Fundação para a Ciência e a Tecnologia (FCT) within the Portuguese Ministry for Science and Education (SFRH/BD/77043/2011).

